# *tailfindr*: Alignment-free poly(A) length measurement for Oxford Nanopore RNA and DNA sequencing

**DOI:** 10.1101/588343

**Authors:** Maximilian Krause, Adnan M. Niazi, Kornel Labun, Yamila N. Torres Cleuren, Florian S. Müller, Eivind Valen

## Abstract

Polyadenylation at the 3’-end is a major regulator of messenger RNA and its length is known to affect nuclear export, stability and translation, among others. Only recently, strategies have emerged that allow for genome-wide poly(A) length assessment. These methods identify genes connected to poly(A) tail measurements indirectly by short-read alignment to genetic 3’-ends. Concurrently Oxford Nanopore Technologies (ONT) established full-length isoform RNA sequencing containing the entire poly(A) tail. However, assessing poly(A) length through basecalling has so far not been possible due the inability to resolve long homopolymeric stretches in ONT sequencing.

Here we present *tailfindr*, an R package to estimate poly(A) tail length on ONT long-read sequencing data. *tailfindr* operates on unaligned, basecalled data. It measures poly(A) tail length from both native RNA and DNA sequencing, which makes poly(A) tail studies by full-length cDNA approaches possible for the first time. We assess *tailfindr’s* performance across different poly(A) lengths, demonstrating that *tailfindr* is a versatile tool providing poly(A) tail estimates across a wide range of sequencing conditions.

## 1 INTRODUCTION

The poly(A) tail is a homopolymeric stretch of adenosines at the 3’-end of the majority of eukaryotic mRNAs. These tails are necessary for the nuclear export of mature mRNAs [1–3] and influence mRNA stability and translation [4]. The poly(A) tail is generated directly after transcription by the non-templated addition of adenosines to the mRNA 3’-end, a process catalyzed by nuclear Poly(A)-polymerases (reviewed in [5]). The initial length of poly(A) tails generated by this process has been proposed to be around 250 nt [6–9]. After nuclear export, poly(A) length is dynamically regulated by the interplay of 3’-to-5’ degradation through exoribonucleases, poly(A) tail stabilization via poly(A) tail binding proteins, and elongation by cytoplasmic Poly(A)-polymerases [10–14]. While it has been shown that the poly(A) tail has a regulatory role, it is still not fully understood whether a specific length allows for specific regulatory outcomes [15]. A minimal poly(A) tail is needed to prevent quick 3’-to-5’ exonuclease degradation [16], yet hyper-adenylated RNAs are marked for fast RNA degradation in the nucleus [15, 17]. Besides from regulating RNA degradation, poly(A) tail length has been shown to correlate with translation efficiency during embryonic development [18, 19], possibly by favoring a closed-loop structure of the mRNA. However, recent studies using *C. elegans* have proposed that shorter poly(A) tails are more actively translated, while longer tails are refractory to translation [20].

To understand the regulatory role of poly(A) tails, it is crucial to be able to measure poly(A) tail length genome-wide with transcript isoform resolution. Up until recently, estimating poly(A) tail lengths was restricted to transcript-specific measurements that relied on PCR and/or on laborious Northern Blotting techniques [21]. These techniques suffer from low throughput, high workload and possible technical artefacts due to amplification [22–24]. Only recently, a set of studies implemented short-read sequencing strategies to study poly(A) tail length in a transcriptome-wide manner [19, 20, 25–28]. While these studies allowed thorough understanding of poly(A) tail lengths throughout the transcriptome for the first time, they are technically restricted to a specific size of poly(A) tails depending on sample enrichment and sequencing strategy. Additionally, most of these techniques rely on PCR amplification of the poly(A) tail region, which might lead to amplification artefacts that affect poly(A) length measurements as well as quantitative comparisons between long and short poly(A) tails [22–24]. Finally, and more importantly, these techniques can only indirectly identify the transcript linked to the poly(A) by alignment of short sequences representing the RNA 3’-ends. Thus it is challenging and in many cases virtually impossible to assign poly(A) tail measurements to specific transcript isoforms.

Oxford Nanopore Technologies (ONT) native RNA Sequencing strategy allows for the sequencing of full-length mRNA molecules without amplification artefacts [29]. The standard library preparation protocol retains the full poly(A) tail in the molecule to be sequenced, making it possible to obtain isoform-specific poly(A) tail length estimates in a transcriptome-wide manner [30]. However, current basecallers do not perform well on long homopolymer DNA regions resulting in the length of poly(A) tails not being accurately reported [31].

Here we present *tailfindr*, an R tool that estimates poly(A) tail length from individual reads directly from ONT FAST5 raw data. *tailfindr* is able to estimate poly(A) tails from both RNA and DNA reads, including DNA reverse-complement reads containing poly(T) stretches. *tailfindr* uses the raw data without prior alignment as input, and estimates the length based on normalisation with the read-specific nucleotide translocation rate. We validate the performance of *tailfindr* on a set of RNA and DNA molecules with defined poly(A) tail lengths. *tailfindr* operates the output of widely used as well as the most recent ONT basecalling applications (Flip-flop model).

## 2 RESULTS

### 2.1 *tailfindr* estimates poly(A) tail length from basecalled ONT native RNA sequencing

Oxford Nanopore Technologies (ONT) Sequencing allows for the sequencing of full-length native RNA molecules containing the entire poly(A) tail by ligation of a double-stranded DNA adapter to the 3’-end of each RNA molecule (Fig. 1A, [30]). Indeed, long stretches of monotonous low-variance raw signal corresponding to poly(A) tails can be observed at the beginning of most reads (Fig. 1B). However, since basecalling relies on fluctuations of the raw signal, these low-variance sections are poorly decoded into the correct nucleobase sequence [31].

**FIGURE 1.**
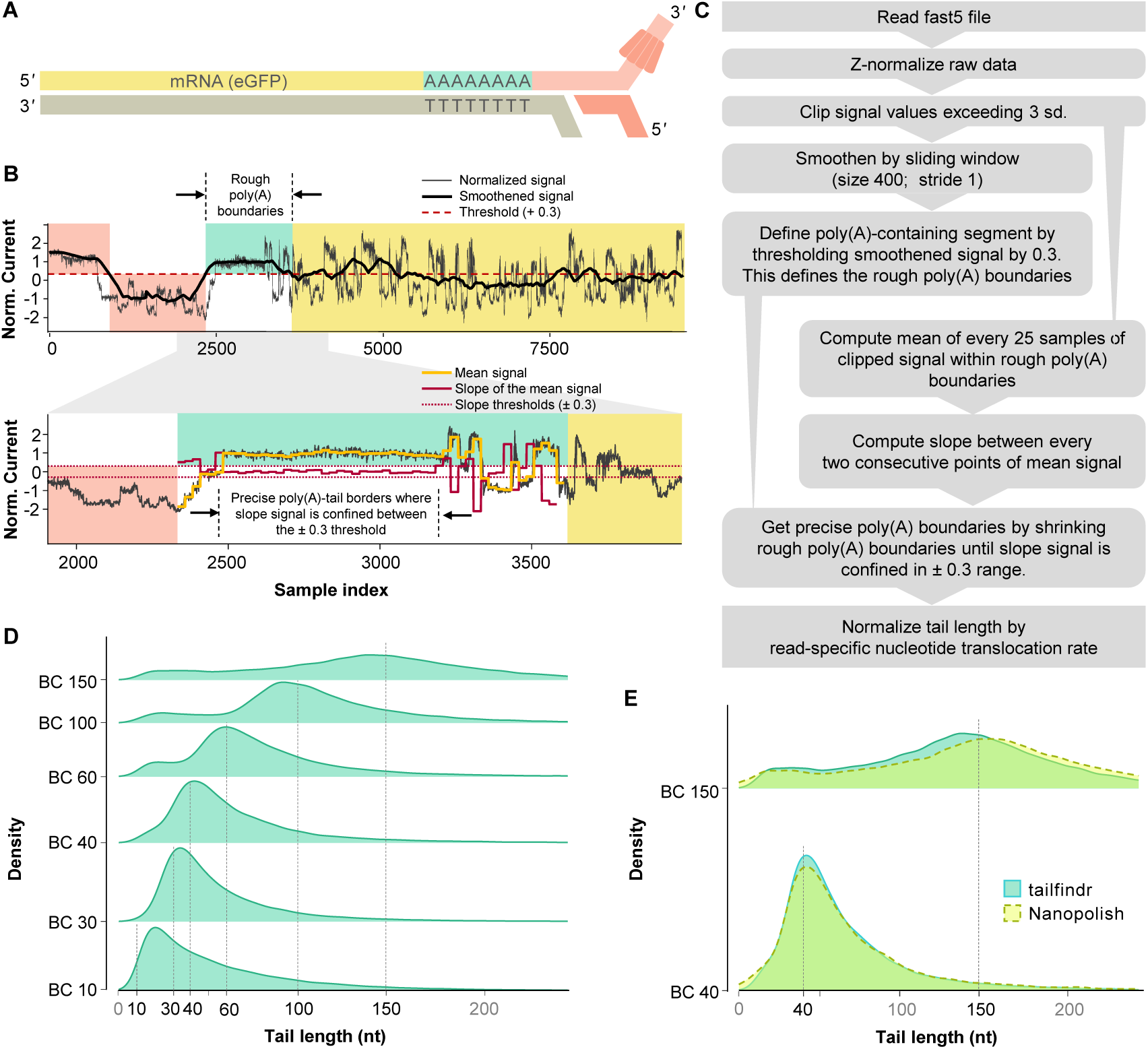
Workflow and performance of *tailfindr* on ONT RNA data. **1A** Schematic representation of Oxford Nanopore RNA Sequencing. The Motor protein (red) is attached to the native RNA molecule (yellow) at the 3’-end by T4 DNA ligation via a double-stranded adapter (light red) with oligo-T overhang. The motor protein thus feeds the RNA strand to the pore from 3’ to 5’. **1B** Representative normalised signal data from eGFP-RNA sequencing. Red background indicates ONT adapter signal, green background represents rough borders of poly(A) signal, yellow background highlights signal from RNA sequence. Boundaries are defined based on *tailfindr* algorithm. **1C** Schematic workflow of data processing by the *tailfindr* algorithm for ONT native RNA sequencing data. **1D** Density plot of poly(A) length estimation on *in vitro* transcribed eGFP-RNA molecules with known poly(A) tail length. Vertical black lines demarcate expected poly(A) length for individual barcodes. **1E** Density plot of poly(A) length estimation from *tailfindr* (turquoise) and Nanopolish (yellow, dashed line) on *in vitro* transcribed eGFP-RNA with poly(A) length of 40 nt or 150 nt.

To identify the region corresponding to the expected poly(A) tail, we apply thresholding to normalized raw data, refine the boundaries of possible poly(A) stretches based on raw signal slope, and normalise by the read-specific nucleotide translocation rate (Fig. 1C, for more details see Materials and Methods). *tailfindr* provides the user with a tabular output containing the unique read-ID, the estimated poly(A) tail length and all factors extracted from the raw data that are needed to calculate the poly(A) tail estimate (Fig. S1A). This allows the user for custom filtering of the acquired poly(A) measurements. Optionally, *tailfindr* allows the user to generate read-specific plots displaying the raw data and all signal derivatives generated in the process to estimate poly(A) tail length (Fig. S1B). To test the performance of our algorithm, we pooled six barcoded *in vitro* transcribed eGFP RNA samples with different poly(A) tail lengths (10 nt; 30 nt; 40 nt; 60 nt; 100 nt; 150 nt) and sequenced the pooled samples with ONT’s native RNA sequencing kit in two replicates. Only reads that cover the full RNA molecule were considered for the analysis. After barcode demultiplexing, the estimated poly(A) tail length match in general with the expected value, with the exception of eGFP with poly(A) tail length of 10 nt (Fig. 1D). While the molecule with expected 10 nt poly(A) length was measured with a mode of 22, the mode of all other barcoded RNA molecules matches well with the expected poly(A) length values (30 nt: 34; 40 nt: 41; 60 nt: 59; 100 nt: 93; 150 nt: 141). However, even though the majority of sequences show the expected poly(A) tail length, the standard deviation of poly(A) tail measurements is rather high (coefficient of variation of around 50%). Thus, while the precision of poly(A) estimation is limited mainly due to outliers towards longer poly(A) lengths (Fig. 1D), the length of most barcoded molecules can be successfully estimated by the use of *tailfindr* on ONT RNA sequencing.

While this study was in progress, another tool estimating poly(A) tail lengths from ONT RNA data has been developed [32]. Instead of estimating poly(A) tails from basecalled data directly, this tool requires read alignment information for the definition of the poly(A) tail segment. To compare whether our algorithm results in similar performance, we measured poly(A) tail lengths from Nanopolish and *tailfindr* on different barcoded eGFP molecules. Our analysis shows that both tools match in precision and length estimation, as exemplified in Figure 1E for 40 and 100 nt poly(A) tail length. However, while both tools agree in the majority of cases in the definition of poly(A) segments, we routinely observed slightly higher estimates from Nanopolish which can be attributed to differences in normalisation (Fig. S2A, S2B). In conclusion tailfindr accurately defines poly(A) tail segments in ONT native RNA sequencing data and provides similar estimates to Nanopolish while only using basecalled data files as input.

### 2.2 Poly(A) and poly(T) tail length can be estimated from ONT DNA sequencing data

ONT native RNA sequencing is lower in both quantity and quality compared to cDNA sequencing approaches and relies on large amounts of starting material (500 ng of poly(A)-selected RNA, [33, 34]). Therefore, cDNA sequencing approaches that retain the full-length poly(A) tail would enable studies where material is scarce as well as increase statistical power of poly(A) tail estimates. We thus aimed to expand tailfindr to operate on ONT DNA sequencing approaches as well. Since standard cDNA approaches result in double-stranded DNA, both poly(A) as well as poly(T) stretches are present in ONT sequencing reads. During cDNA sequencing both of these strands are threaded through the pore separately from 5’ to 3’ (Fig. 2A). Indeed we observe homogenous stretches of raw signal both at the beginning (poly(T) tail) as well as at the end (poly(A) tail) of individual raw read sequences (example for poly(T)-containing read in Figure S3B).

**FIGURE 2.**
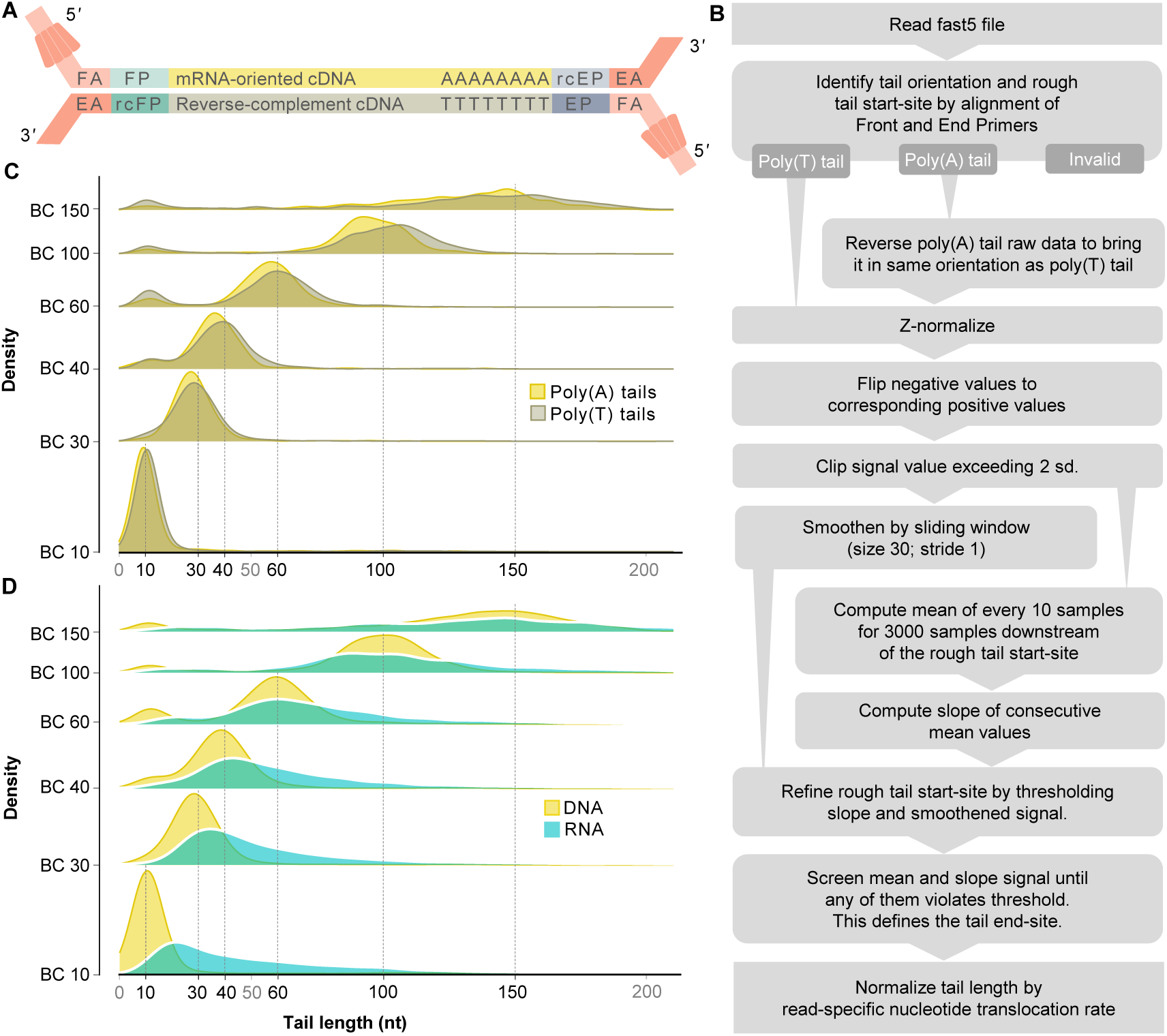
Workflow and performance of *tailfindr* on ONT DNA sequencing data. **2A** Schematic representation of Oxford Nanopore DNA Sequencing. In cDNA approaches, amplification is ensured by oligo-dT-aided anchoring of the End Primer (EP, blue) and addition of Front Primer sequence (FP, green) by strand-switch during reverse transcription. The Motor protein (red) is attached to the double-stranded DNA molecules at both ends by T4 DNA ligation. The Front Adapter (FA) bears the motor protein, while the End Adapter (EA) is a short complementary oligo that will ultimately appear at the 3’-end of resulting sequences. Both DNA strands will be sequenced from 5’ to 3’. Thus, oligo-dT stretches will be present at the beginning of raw data, while oligo-dA stretches appear at the end. **2B** Schematic workflow for ONT DNA sequencing data processing by the *tailfindr* algorithm. **2C** Density plot of poly(A) (yellow) and poly(T) (grey) length estimates on PCR-amplified eGFP coding sequence with known poly(A) length. Vertical black lines demarcate expected poly(A) length for individual barcodes. **2D** Density plot of poly(A) length estimates on RNA (turquoise) vs poly(A)/(T) length measured on DNA sequences (yellow).

We extended our algorithm to accommodate ONT DNA sequencing data output (Fig. 2B). Most significantly, we account for the double-stranded nature of DNA and define the read type (poly(A)- or poly(T)-containing) by making use of known sequence motifs in Nanopore Adapters (details in Material and Methods). As for RNA sequencing, *tailfindr* provides the user with a tabular output of tail length measurements (including the read type) as well as optional raw data plots (Figure S3). We tested the performance of the DNA-specific *tailfindr* algorithm on PCR products of eGFP coding sequence with known poly(A)/(T) length in two replicates, similar to the spike-ins generated for native RNA sequencing. As shown in Figure 2C, the DNA-specific *tailfindr* approach resulted in estimated poly(A) and poly(T) lengths close to the expected length for barcoded molecules (mode of distribution for 10 nt: 10; 30 nt: 28; 40 nt: 38; 60 nt: 59; 100 nt: 92 for poly(A) and 110 for poly(T); 150 nt: 148 for poly(A) and 155 for poly(T)). For all poly(A)/(T) tail lengths bigger than 10 nt, a small subpopulation of reads with shorter estimated tails could be observed, possibly due to incorrectly assigned barcodes. While poly(A) and poly(T) estimates show very similar density distributions for length estimates, poly(T) lengths are routinely slightly longer than estimates for poly(A) lengths.

Next we compared poly(A)-length estimates from PCR-DNA and native RNA sequencing. We observed that DNA sequencing results in significantly more precise estimation of poly(A) tail length, mainly due to fewer outliers towards longer poly(A) tail lengths (Fig. 2D). Especially the shortest poly(A) tail length (10 nt) could be estimated more correctly in DNA sequencing (mode of poly(A) length estimation 10 in DNA vs 22 in RNA sequencing). On other poly(A) lengths the mode of poly(A) estimation does not differ dramatically, but the precision of measurement is significantly higher in DNA sequencing approaches (coefficient of variation 50% in native RNA sequencing vs 30% in DNA sequencing). In summary, *tailfindr* is able to estimate poly(A) and poly(T) tail size from ONT DNA sequencing with significantly higher precision compared to ONT RNA sequencing estimation.

### 2.3 *tailfindr* is compatible with Flip-flop model basecalling

While this manuscript was in preparation, ONT released a new DNA basecalling strategy based on Flip-flop models. Flip-flop model basecalling screens the raw signal by comparing probabilities to either stay in the same nucleotide state or change to a new state. Additionally, the raw data is read by averaging over two sample points, as opposed to averaging over five sample points in standard model basecalling. These improvements have been shown to result in higher quality basecalling, and more importantly to increase the basecall fidelity over homopolymer sequences [35]. So far, Flip-flop model basecalling is only available for ONT DNA sequencing data.

We implement changes in *tailfindr* to account for the updates in Flip-flop model raw data output. As expected, Flip-flop model basecalling detects more nucleotide translocations (called ‘moves’) over poly(A) stretches when compared to standard model basecalling (Fig. 3A, yellow highlights). To test whether the detected moves agree with expected poly(A)/(T) length, we plotted the moves from either standard model basecalling (Fig. 3B) or Flip-flop model basecalling (Fig. 3C) on eGFP-PCR products with 30 nt or 100 nt poly(A)/(T) tail length. While Flip-flop model basecalling resulted in significantly more moves over poly(A)/(T) tail sections compared to standard model basecalling, the detected number of moves still severely underestimate the existing poly(A)/(T) lengths. Thus even with improved homopolymer basecalling fidelity, external tools are needed to correctly measure poly(A) tail lengths. We used *tailfindr* to compare poly(A) and poly(T) tail measurements from the same sequencing reads basecalled either with Flip-flop or standard models, and could show that the estimated poly(A)/(T) tail length is highly correlated between the two basecalling approaches (R = 0.93 for poly(A); R = 0.97 for poly(T); Fig. 3D,E). We thus conclude that *tailfindr* operates on both standard and the most recent Flip-flop model basecalling, and provides accurate poly(A)/(T) length estimates for ONT DNA sequencing approaches.

**FIGURE 3.**
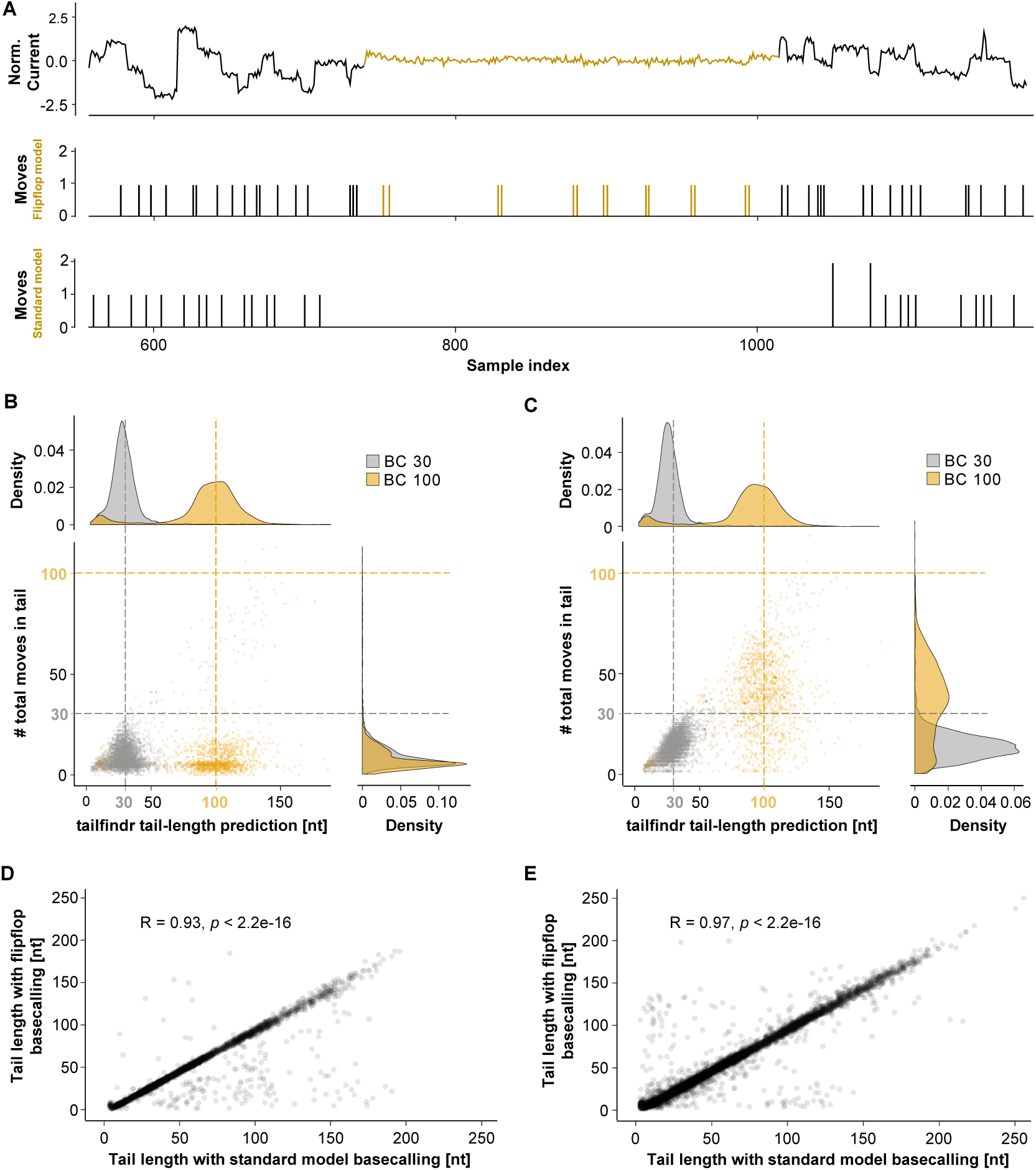
Differences in poly(A) tail estimation for standard and Flip-flop model basecalling. **3A** Representative raw data squiggle of PCR-amplified eGFP coding sequence over the identified poly(A) tail region (colored yellow) with associated moves (shifts in raw data representing possible nucleotide translocations) in both Flip-flop (top) and standard (bottom) model basecalling. Flip-flop model basecalling detects moves with higher resolution, and calls more moves especially in the poly(A) tail region (yellow). **3B,C** Scatter plot of estimated poly(A)/(T) tail length (x-axis) and moves detected with standard **(B)** or Flip-flop model basecalling **(C)** on PCR-amplified eGFP coding sequence with poly(A) length of 30 nt (grey) and 100 nt (yellow). Colored dashed lines indicate expected poly(A) length. **3D,E** Scatter plot of poly(A) **(D)** or poly(T) **(E)** tail length estimated from PCR-amplified eGFP coding sequence with different poly(A) tail lengths basecalled either with standard (x-axis) or Flip-flop model (y-axis). (R, p by Pearson correlation)

## 3 DISCUSSION

Polyadenylation at the 3’-end is understood to be a major regulator of mRNA [1–4]. While the poly(A) length of mRNAs has been under investigation since the 1970’s [36–39], transcriptome-wide analysis of poly(A) tail lengths have only recently emerged. The advent of Oxford Nanopore Technologies (ONT) native RNA Sequencing technology now allows direct sequencing of full-length mRNA molecules, which intrinsically contain their full poly(A) tail, unbiased by potential amplification artefacts [29]. However, even the most recent updates in basecalling tools do not perform well over long homopolymeric sequence stretches [31, 35].

In this work we present *tailfindr*, a versatile R tool that allows estimation of poly(A) tail lengths from basecalled ONT long-read sequencing data from both native RNA and DNA sequencing approaches. We show that *tailfindr* is able to detect the poly(A) tail boundaries of *in vitro* transcribed eGFP RNA molecules and estimate their lengths based on read-specific raw data normalisation. For molecules with known poly(A) tails from 30 nt up to 150 nt the estimates match well with the expected lengths (Fig. 1D), however the shortest poly(A) tail (10 nt) was estimated to have longer tails than expected. We believe that this bias can be explained by sample contamination of this RNA molecule during preparations, or by inefficient oligo-dT sequencing adapter ligation to poly(A) tail stretches at or below 10 nt. Consistent with the latter explanation we observed that the barcoded 10 nt RNA molecule was underrepresented in the RNA sequencing libraries compared to input quantities (data not shown). Overall, *tailfindr* correctly estimates poly(A) tail lengths of *in vitro* transcribed RNAs over a wide range of lengths.

We further show that *tailfindr* poly(A) tail estimates agree closely with a recently developed tool that relies on prior mapping of the data [32]. While poly(A) tail boundaries in raw signal are found to be essentially the same with the two different approaches (Fig. S2C,D), the final calculated poly(A) tail lengths differ (Fig. S2A). Specifically, *tailfindr* estimates short poly(A) stretches slightly longer than Nanopolish, while long poly(A) stretches result in shorter estimates in *tailfindr*. These differences can be explained by different calculation of the average nucleotide translocation rate (Fig. S2B) which is used to normalise raw poly(A) tail measurements. Nanopolish normalises by calculating the read-specific median of the samples per nucleotide after removing 5% of the translocation rate outliers. We have observed that this normalisation is resulting in correct poly(A) estimation in RNA, but not DNA sequencing approaches.

Instead we normalise by the read-specific geometric mean of samples per nucleotide without outlier removal. Another difference between the tools is that tailfindr does not need any sequence data preprocessing, as it only requires basecalled FAST5 files as input. This allows for poly(A) tail studies independent of any other tool than the essential basecaller, which would allow for an integration of the tailfindr algorithm into the basecalling procedure. This in turn makes it possible to assign poly(A) tail lengths to individual reads in parallel to the sequencing procedure, making live poly(A) tail analysis feasible.

In comparison to recent short-read sequencing-based strategies to measure poly(A) tails, methods using ONT sequencing are currently less precise. Short-read sequencing approaches promise poly(A) measurements with just a few bases of deviation due to cyclic incorporation of nucleotides and integration of the fluorescence signal of multiple molecules towards one single basecall [20, 25–28]. In contrast, ONT long-read sequencing measures individual single-stranded molecules, and single nucleotide changes are detected based on subtle changes in measured current levels. More importantly, the raw signal for ONT sequencing does not change over a homopolymeric region, making single-event detection almost impossible. Thus, ONT poly(A) length estimation relies on normalisation rather variable data taken from single-molecule measurements. Most of the variation observed in *tailfindr* poly(A) estimation thus comes from the sequencing process. However, the sequencing chemistry as well as the properties of the motor protein are under constant development. It is thus conceivable that in the near future an increase in speed and robustness of translocation rates can be observed, which will have a positive impact on poly(A) tail estimation [35].

While not being as precise in measuring poly(A) tails, ONT long-read sequencing approaches have unique advantages over short-read sequencing approaches. First, ONT sequencing is intrinsically a single-molecule technique. Second, RNA sequencing approaches are amplification-free, avoiding the emergence of possible amplification artefacts. Third, since the native molecule is sequenced as it comes from the specimen, additional features of the RNA can measured directly, as was shown for RNA modifications [32, 40]. Fourth, and most importantly, long-read sequencing allows direct assignment of transcript isoforms to single molecules without bioinformatics post-processing, making truly isoform-specific measurements of poly(A) tail lengths possible. Additionally, ONT sequencing allows to study features of 5’-end and 3’-end events of the same molecule in conjunction with the poly(A) tail length. Together, ONT sequencing in conjunction with *tailfindr* poly(A) estimation offers great potential to combine the study of poly(A) tail length and other RNA features with transcript-isoform specificity in one assay.

Next to handle data from ONT RNA sequencing applications, *tailfindr* is the first tool to show that poly(A) tails can be measured in ONT DNA sequencing. We further modified *tailfindr* to handle the most up-to-date basecalling approaches with Flip-flop models. We could show that poly(A) tail measurements from ONT DNA sequencing are more precise compared to measurements of similar RNA molecules (Fig. 2D). This is mainly explained by a faster and more robust translocation rate with less likelihood for stochastic stalling during sequencing. Interestingly, poly(T) stretches are commonly estimated slightly longer than poly(A) stretches from the same sample (Fig. 2C). A possible explanation for this would be a slightly faster translocation speed of purine stretches compared to pyrimidine stretches, and could be accounted for in more specific normalisation strategies in the future.

The existence of *tailfindr* makes it possible to design specific cDNA library preparation protocols that retain the full poly(A) tail in ONT sequencing approaches. This strategy has recently been shown to allow further insights into poly(A) tail regulation based on PacBio long-read sequencing [41]. ONT cDNA sequencing has the advantage to yield approximately 10x more data per library preparation compared to native RNA sequencing, and due to amplification would allow sequencing experiments starting with minute RNA amounts as input [34, 42]. Additionally, we envision that future cDNA applications that include Unique Molecular Identifiers (UMI) will make it possible to acquire multiple poly(A) tail measurements for each molecule, which will increase the fidelity of isoform-specific poly(A) tail measurements. Thus, using *tailfindr* with specific ONT cDNA applications offers new approaches to study the role of poly(A) tail lengths from scarce biological samples.

In conclusion, ONT RNA sequencing offers a new possibility to study poly(A) tail biology, especially by directly associating poly(A) tail length with other RNA features in a transcript isoform-specific manner. *tailfindr* has proven successful in measuring the poly(A) tail of both RNA and DNA sequencing solely from basecalled raw data, an approach that allows real-time analysis during ONT sequencing. With the application of *tailfindr* for ONT DNA sequencing we allow future development of poly(A)-retaining cDNA sequencing assays that further increase the ability to study poly(A) tail lengths of scarce material.

## 4 MATERIALS AND METHODS

### 4.1 Spike-in generation

To generate RNA with known poly(A) tail lengths, we used eGFP as a carrier RNA as it fulfils basic criteria for successful ONT RNA sequencing (especially minimal length requirement). The coding sequence of eGFP was amplified from pCS2+-eGFP vector using High Fidelity Phusion MasterMix (ThermoFisher, #F-531L). The primers for the PCR included the SP6 promoter sequence and a barcode in the forward primer, as well as the a homopolymer T stretch in the reverse Primer (see Table 1). After gel-purification of the desired PCR product, a second PCR was performed with a reverse Primer that introduces a Bfo1 restriction site before the homopolymer T stretch (polyA Bfo1 rev, together with SP6 Bfo1 fw, Table 1). After gel-purification and Phenol-chloroform extraction, the resulting PCR products were used for Nanopore DNA Ligation Sequencing (see below). For preparation of RNA spike-ins, the PCR products were digested with FastDigest Bfo1 (ThermoFisher, #FD2184) for 2 hours and purified by Phenol-chloroform extraction. 100-300 ng of purified DNA were used for RNA *in vitro* transcription by the SP6 mMessage mMachine kit (ThermoFisher, #AM1340) following manufacturer’s procedures. The resulting RNA was purified using Zymo RNA Clean & Concentrator-5 columns (Zymo Research, #R1013).

**TABLE 1.**
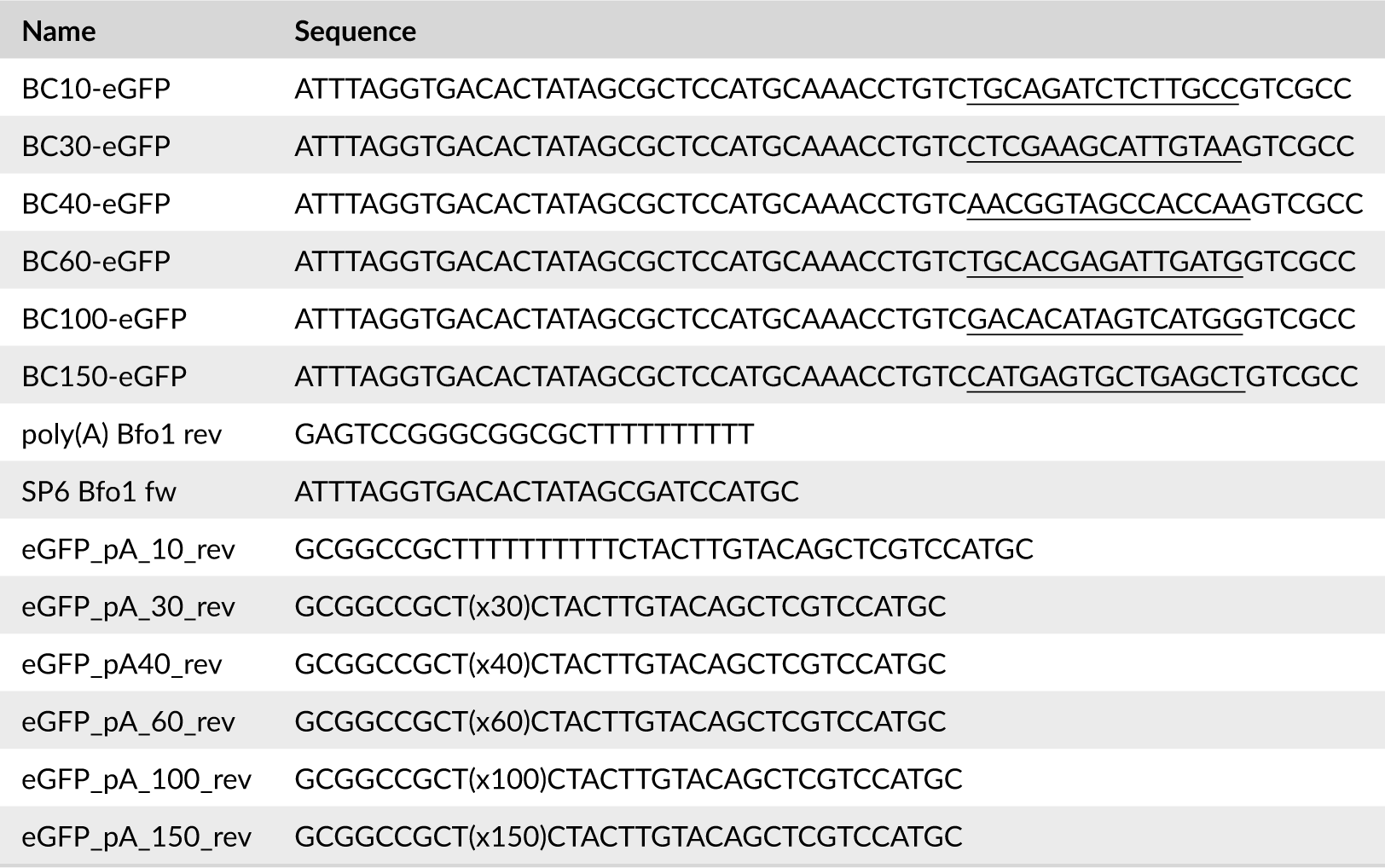
DNA oligos for the design of poly(A)-tailed eGFP constructs.

### 4.2 ONT Long-read Sequencing

Native RNA Sequencing was performed using the ONT kit SQK-RNA001 following the manufacturer’s protocol. In brief, 500 ng of poly(A)-selected RNA were mixed with 100 ng of poly(A) spike-in RNA, or 500 ng poly(A) spike-in RNA were used alone. The RNA was ligated to ONT RT adapter (RTA) and used for Reverse Transcription with SuperScript II (ThermoFisher, #18064022). Next, the proprietary Sequencing Adapter was ligated using T4 DNA ligase (NEB, #M0202M) and loaded onto ONT Sequencing Flow Cells (FLO-MIN106 R9.4.1). Sequencing was performed for 16-24 hours using MinKNOW 2 software. All RNA purification steps were performed with RNAClean XP beads (Beckham Coulter, #A63987) with 15 minutes incubation intervals. DNA Sequencing was performed using the DNA Ligation Kit SQK-LSK108 on poly(A) containing PCR products. In brief, 500 ng of pooled barcoded PCR products were end-prepped using the NEBNext Ultra II dA tailing module (NEB, #E7546S) and ligated to proprietary Sequencing Adapters using T4 DNA ligase (NEB, #M0202M). Purified libraries were sequenced on Flow Cells (FLO-MIN106 R9.4.1) for 24 hours using MinKNOW 2.

### 4.3 Sequencing data processing

RNA and DNA raw reads were basecalled using Albacore v2.3.3. DNA raw reads were additionally basecalled with Guppy v2.3.1 using Flip-flop models. Sequencing quality and general metrics were assessed using NanoPlot (v1.19.0, [43]). Reads that passed the default albacore quality filter were mapped against the eGFP sequence using minimap2 (v2.14-r883) with default settings for ONT data mapping (-ax splice -uf −14 for RNA; -ax splice for DNA, [44]).

### 4.4 Demultiplexing barcoded spike-ins

All alignments discussed in this manuscript, unless mentioned otherwise, were performed using Smith-Waterman local alignments with Biostrings (match score 1; mismatch score −1; gap opening penalty 0; and gap extension penalty 1) [45]. The normalised alignment score was calculated by dividing the local alignment score by the length of the query sequence. If not otherwise mentioned, alignments with a normalised alignment score below 0.6 were discarded as unspecific.

Barcoded eGFP RNA reads with known poly(A) length were demultiplexed by locating the first 29 bases of eGFP sequence (see Table 2) within the first 250 bases of FASTA strings extracted from every FAST5 file. Next, the barcode was assigned by aligning the expected barcode sequences against the extracted read sequence preceding the eGFP alignment (see Table 2). The barcode with highest normalised alignment score (and above threshold of 0.6) was assigned to the read. To analyze barcoded eGFP DNA reads, the orientation of reads was investigated by aligning the first 29 bases of eGFP and its reverse-complement (Table 2) to the first 250 bases of FASTA strings extracted from each FAST5 file. A read was considered a poly(A)-containing read if the normalised alignment score of eGFP sequence was greater than both the normalised alignment score of the reverse-complement of eGFP and the threshold value of 0.5. Reads where the normalised alignment score of the reverse-complement of eGFP was higher than the forward eGFP sequence and passed the threshold value of 0.5 were considered to be poly(T)-containing reads. For Barcode demultiplexing, first the sequence preceding the identified eGFP start was queried for the presence of the experiment-specific PCR Front Primer in case of poly(A) reads, or its reverse-complement for poly(T) reads (sequences in Table 2). Next, the sequence between Front Primer and eGFP locations was used for barcode identification as described above.

**TABLE 2.**
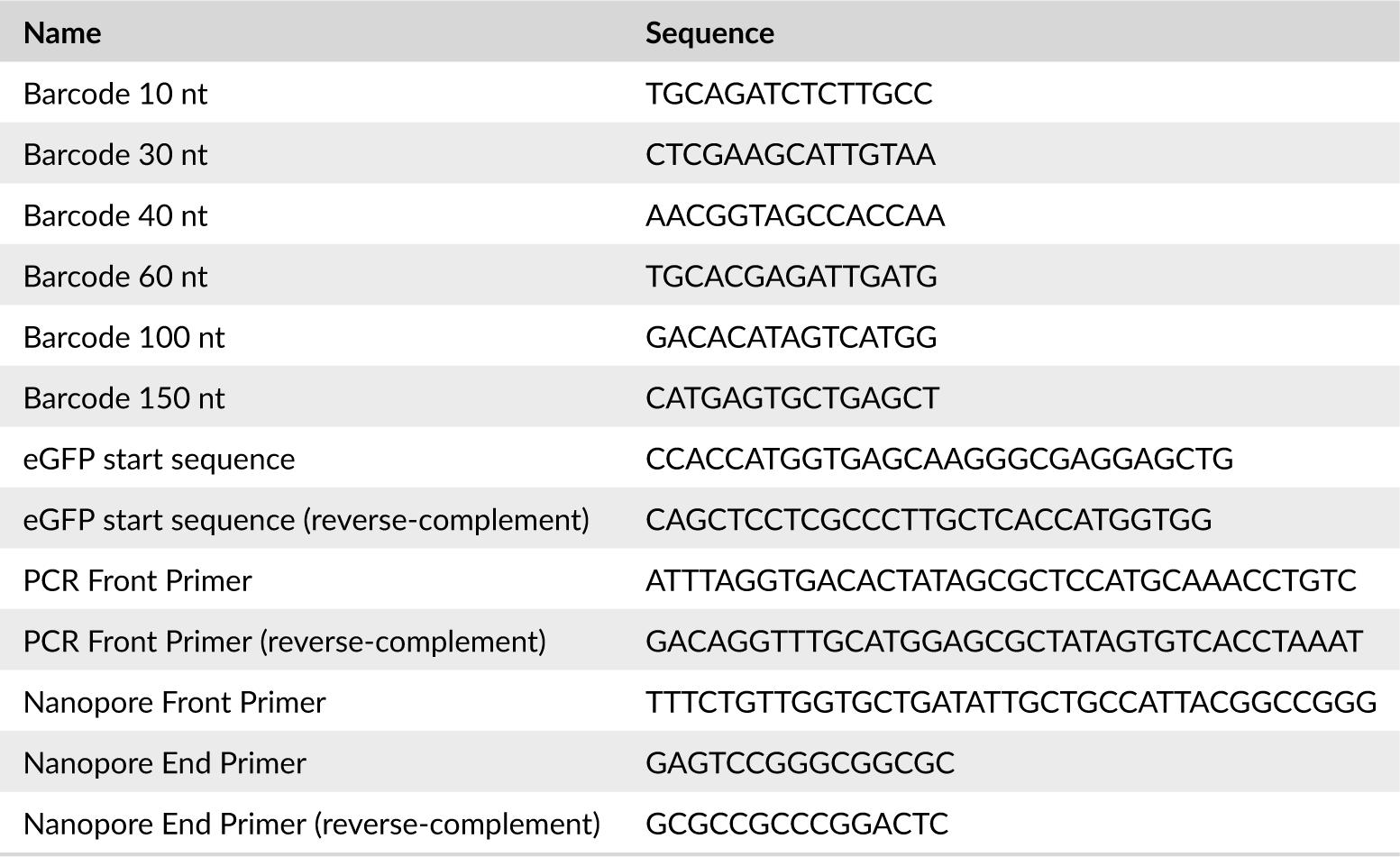
Sequences used in *tailfindr* alignments.

### 4.5 *tailfindr* RNA poly(A) length estimation algorithm

To identify the signal corresponding to the poly(A) tails in RNA reads, the raw signal from ONT native RNA sequencing is extracted from the FAST5 files and z-normalised. Next, signal values above +3 and below −3 are truncated. The resulting processed raw signal is smoothened by a moving average filter (window size 400 samples; stride 1) in both directions separately. Both smoothened signal vectors are then merged by point-by-point maximum calculation. Next, the calculated smoothened signal is segmented into regions being above or below 0.3. The expected signal of the ONT adapter consists of one segment above and one segment below 0.3 in smoothened signal. The poly(A) tail immediately follows the Nanopore Adapter, thus the next segment in which the smoothened signal is above 0.3 is considered the poly(A) region, and the boundaries of this segment are considered the rough start and end of poly(A) tail (Fig. 1B).

The rough start and end are refined by first calculating a mean signal of the processed raw data contained between these boundaries through a moving average filter (window size 25; stride 25). Next, the slope of this mean signal is calculated between each two consecutive points. The boundaries of the longest continuous stretch of low-slope values (confined within bounds of +0.3 and - 0.3 of slope signal) between the rough poly(A) start and end boundaries are considered the precise boundaries (Fig. 1B). The resulting poly(A) tail measurement in sample points is then normalised by the read-specific nucleotide translocation rate (see below).

### 4.6 *tailfindr* DNA poly(A)/(T) estimation algorithm

Unlike RNA, DNA is double-stranded. Thus, homopolymer poly(A) and poly(T) stretches can occur. To determine the read orientation, the Nanopore-specific Front and End Primer sequences (sequences in Table 2) are aligned against the first 100 bases extracted from FAST5 files. A read is considered poly(T)-containing, if the normalised alignment score of End Primer sequence is greater than that of the Front Primer sequence, and above the threshold of 0.6. Conversely, a read is considered poly(A)-containing if the normalised alignment score of Front Primer sequence is greater than that of the End Primer sequence, and above the threshold of 0.6. To ensure that the full poly(A) tail is present in raw data, the last 50 bases of poly(A)-containing reads are queried for the presence of the reverse-complement End Primer sequence. Reads where the normalised alignment score of the reverse-complement End Primer is below 0.6 are considered truncated poly(A) reads and not analysed further.

To identify borders of poly(A) or poly(T) stretches by similar procedures, the raw data of poly(A)-containing reads is reversed. Thus, both the poly(A) and poly(T) stretch are expected to be at the beginning of raw signal. The alignment of End Primer is considered the approximate start of the poly(A) or poly(T) stretch. Next, the raw data is z-normalised and converted to absolute values. To reduce computational workload, calculations to identify precise borders of poly(A)/(T) stretches are restricted to 3000 raw samples downstream of the rough poly(A)/(T) start site. This 3000-samples wide search window is wide enough to accomodate poly(A)/(T) tails of approximately 350 nt length. The mean signal is generated by applying a sliding window (window size 10; stride 10) to the processed raw signal. Next, the slope of this mean signal is calculated between every two consecutive points. The precise start of the respective tail is considered to be the first location after the rough start site where the calculated slope is between −0.2 and 0.2, and the mean signal is between 0 to 0.3. To identify the precise tail end, the slope and the mean signals downstream of the precise tail start site are tested for violating their respective thresholds (see above). Since short non-tail-like signal spikes can randomly occur, we test the signal downstream of this tentative tail end for tail-like signal within thresholds until we either reach the end of the search window of 3000 sample points, or find another stretch of tail-like signal of at least 60 sample points in length. In the latter case, the tentative tail end is updated to the downstream tail end to account for the spike signal. The maximum allowable signal length exceeding the threshold that is located between two tail-like signal has been to set to 120 nt (e.g. 120x read-specific nucleotide translocation rate).

The difference between the precise boundaries define the raw length of poly(A)/(T) stretches in sample points. This value is normalised by the read-specific nucleotide translocation rate calculated dependent on the respective base-calling strategy (see below).

### 4.7 Calculation of the read-specific nucleotide translocation rate

The translocation speed of the biological molecule through the pore is not homogenous. Thus, the translocation rate for each individual nucleotide can differ significantly. The translocation speed can be influenced by the sequencing context [31], but also by sample time and sequencing buffer conditions (unpublished observations). Furthermore, the translocation speed can be influenced by RNA or DNA modifications, which however should not affect RNA sequencing from *in vitro* transcribed molecules, or DNA sequencing PCR-amplified molecules. Importantly, the motor protein for RNA and DNA differs, leading to dramatically different average translocation speed (70 nt for RNA in ONT Kits SQK-RNA001 vs. 450 nt for DNA in ONT Kits SQK-LSK108). In conclusion it is important to estimate the average nucleotide translocation rate for each read separately to account for the specific conditions at which the read was recorded. In basecalled ONT sequence data, a ‘move’ in raw data describes a single-nucleotide translocation through the pore. To calculate the average read-specific nucleotide translocation rate, we first extract a vector containing the number of sample points per move from the FAST5 events table of each individual read. If a move of 2 is detected (does not occur in basecalling with Flip-flop models), we divide the number of sample points by 2, as we reason that a preceding nucleotide translocation was not detected by the basecaller. From the resulting distribution of sample points per move, we then compute the geometric mean. This strategy results in robust estimation of poly(A) tail length for both ONT RNA and DNA sequencing approaches with standard model basecalling (Fig. 1D; Fig. 2C). In our experience, this approach is resulting in more robust normalisation compared to normalisation by the median of single-nucleotide translocation rates, which is used in Nanopolish (Fig. S2B; [32]).

Oxford Nanopore Technologies recently presented an updated basecalling strategy with Flip-flop models instead of standard models used for neural network nucleotide decoding [35]. In this approach, the raw current level data is decoded by averaging 2 sample points instead of 5, resulting in higher-resolution basecalling. Additionally, we could not observe moves of 2, which in most cases represent missed nucleotide translocations. When we calculate the read-specific nucleotide translocation rate from reads basecalled with standard and Flip-flop models, we routinely observe lower average values for Flip-flop model basecalling. The calculated geometric mean of moves is often below 8. The geometric mean calculated from standard model basecalling routinely results in translocation rates between 8 and 9, which agrees with average translocation rates communicated by ONT (unpublished communications). Compared to standard model basecalling it is possible that Flip-flop model basecalling over-segments the raw data, resulting in too low average nucleotide translocation rates when calculated by the geometric mean. This is exemplified by the observation that the most frequently observed nucleotide translocation rate is 2 samples, the lowest possible value considering the resolution of 2 sample points. Instead of calculating the average rate by geometric mean, we used the arithmetic mean after discarding the 5% highest outliers as an approximation of the right normaliser, based on comparisons to results obtained by standard model basecalling on the same PCR-DNA molecules.

## Acknowledgements

MK would like to acknowledge the constant scientific exchange and troubleshooting with, as well as critical assessment of the manuscript by Kirill Yefimov and Teshome Bizuayehu. MK likes to further thank his family (small and big) for constant moral support during his career.

## Supplementary Materials

**FIGURE S1.**
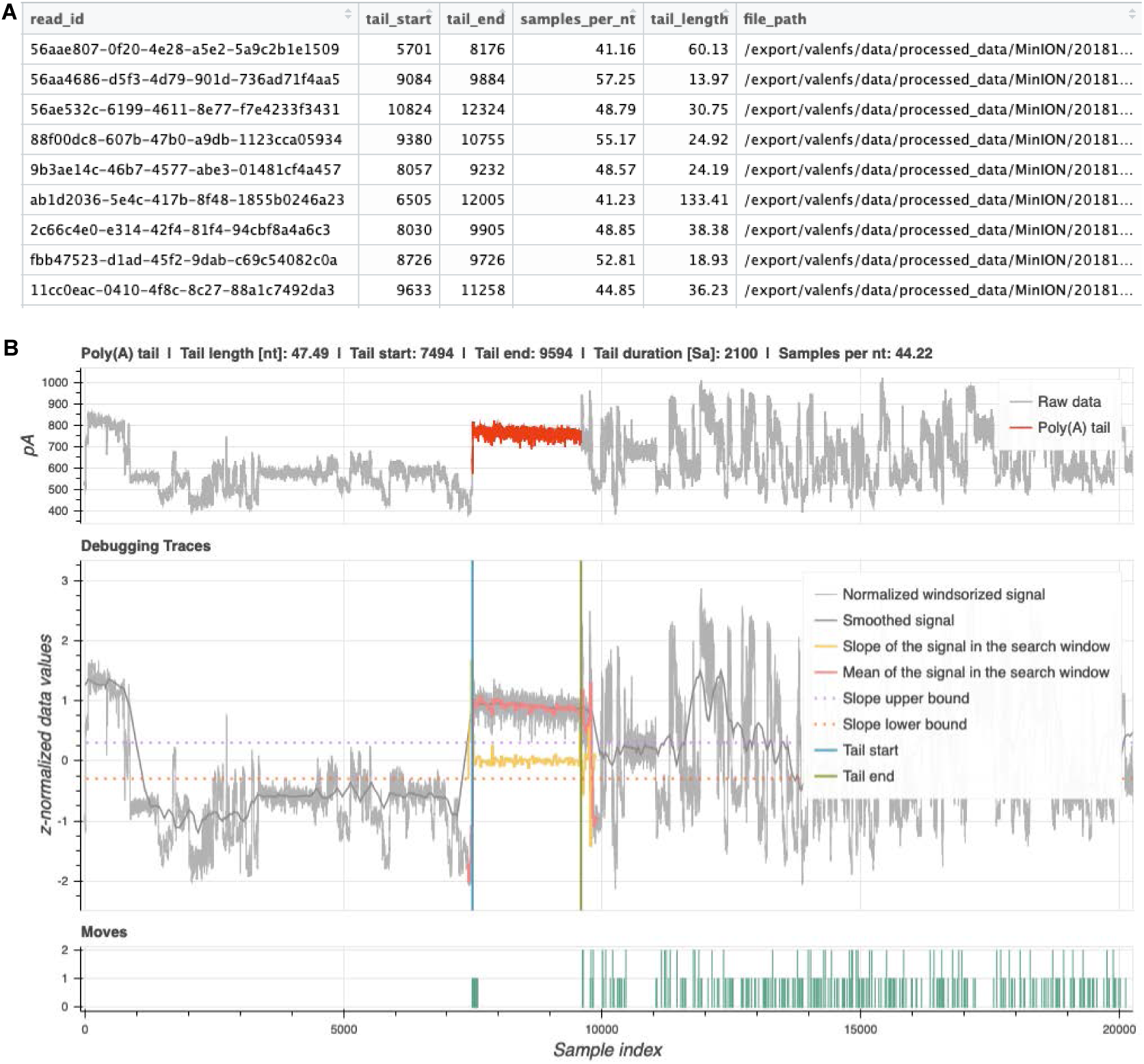
Example data output from *tailfindr* on RNA sequencing data. **S1A** Tabular output containing the unique ONT read ID, tail start and end coordinates, the normaliser (samples_per_nt), the calculated tail length in nt (tail_length) and the filepath for each individual read. **S1B** Example debugging plots that can be optionally generated by *tailfindr* for highlighting the poly(A) tail. Displayed is a poly(A) tail of *in vitro* transcribed eGFP RNA. Top panel shows the total raw data with highlighted poly(A) section (red). Middle panel shows calculated signal derivatives used to define the poly(A) tail boundaries for debugging purposes. Bottom panel displays detected moves as recorded by the basecaller (green).

**FIGURE S2.**
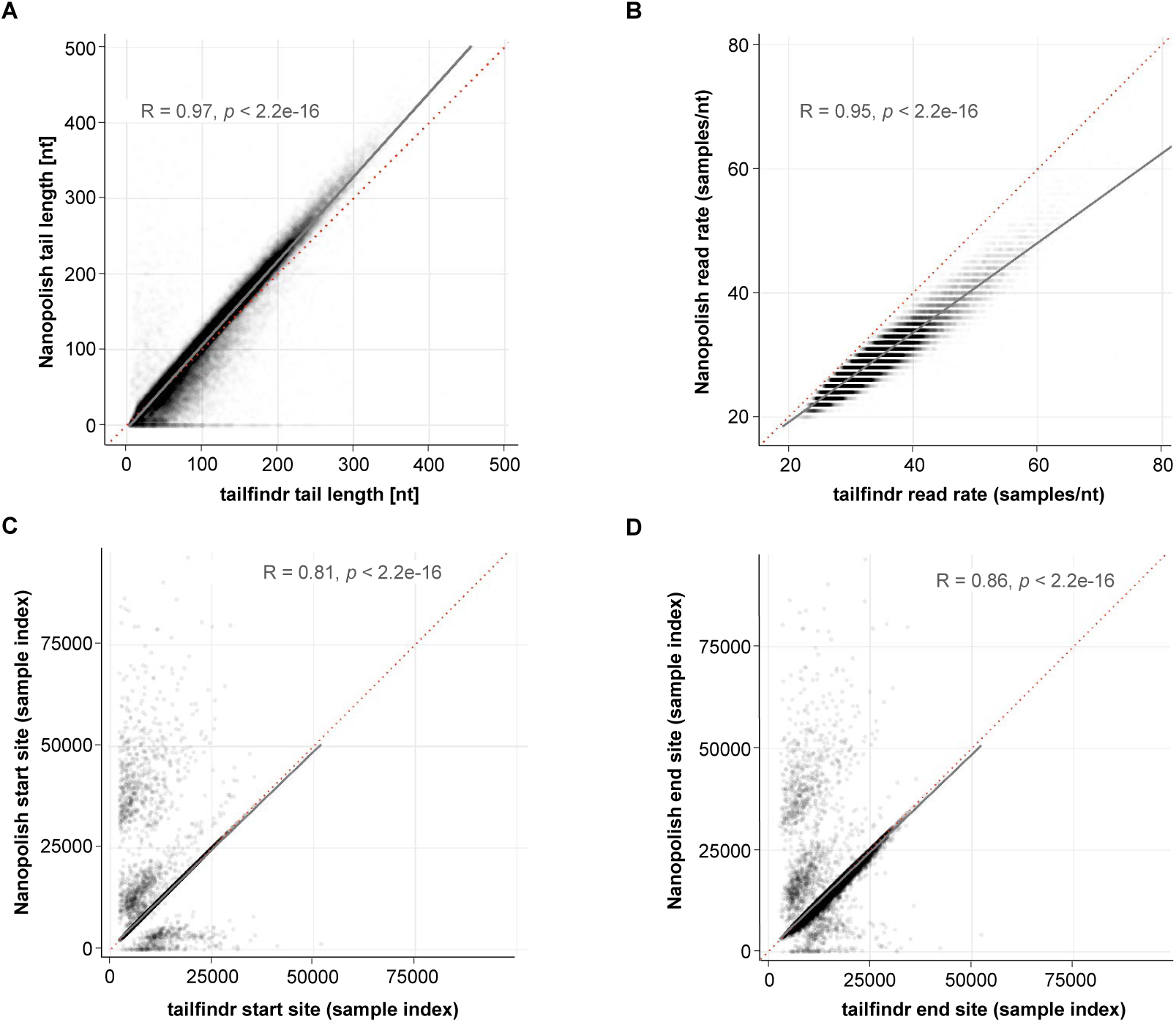
Comparison of poly(A) tail estimation from *tailfindr* and Nanopolish. **S2A** Scatter plot of poly(A) length estimation from *tailfindr* (x-axis) and Nanopolish (y-axis) on *in vitro* transcribed eGFP-RNA with different poly(A) length. Grey line indicates Pearson-correlation fit, red dashed line indicates x=y. (R, p by Pearson correlation) **S2B** Scatter plot of read-specific nucleotide translocation rate used to normalise poly(A) tail length from *tailfindr* (x-axis) and Nanopolish (y-axis). Grey line indicates Pearson-correlation fit, red dashed line indicates x=y. (R, p by Pearson correlation) **S2C** Scatter plot of identified poly(A) start from *tailfindr* (x-axis) and Nanopolish (y-axis). Grey line indicates Pearson-correlation fit, red dashed line indicates x=y. (R, p by Pearson correlation) **S2D** Scatter plot of identified poly(A) end from *tailfindr* (x-axis) and Nanopolish (y-axis). Grey line indicates Pearson-correlation fit, red dashed line indicates x=y. (R, p by Pearson correlation)

**FIGURE S3.**
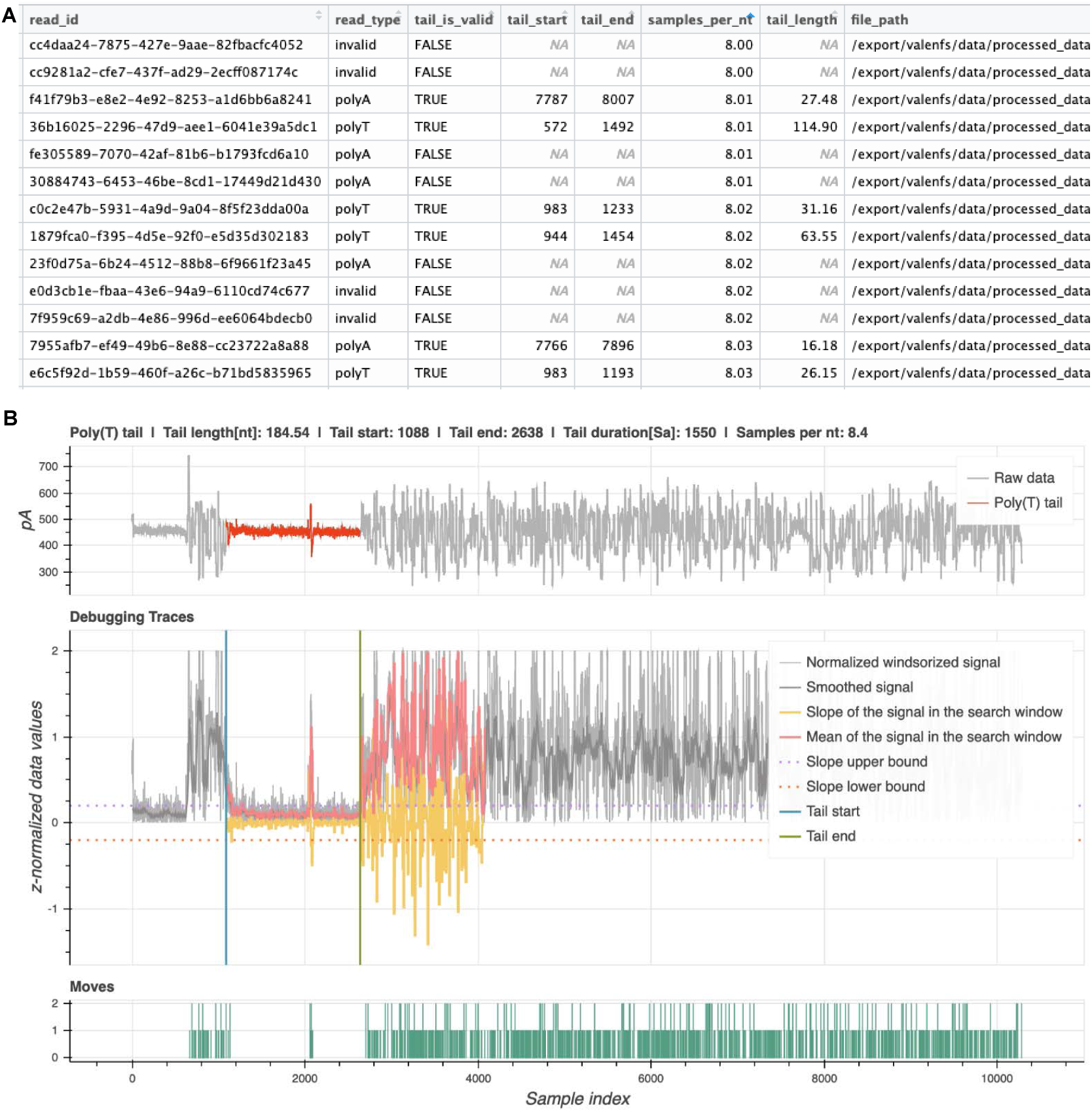
Example data output from *tailfindr* on DNA sequencing data. **S3A** Tabular output containing the unique ONT read ID, read orientation (read_type), tail start and end coordinates, the normaliser (samples_per_nt) and the calculated tail length in nt (tail_length), among others. **S3B** Example debugging plots that can be optionally generated by *tailfindr* for highlighting the poly(A)/(T) sections. Displayed is a poly(T) stretch of eGFP PCR-DNA basecalled with standard model. Top panel shows the total raw data with highlighted poly(T) tail section (red). Middle panel shows calculated signal derivatives used to define the poly(T) tail boundaries for debugging purposes. Bottom panel displays detected moves as recorded by the basecaller (green). Important to notice is a small signal spike in the raw data that *tailfindr* allows to be included in the full poly(T) tail.

